# THE COMPARATIVE STUDY OF THE EFFECT OF LOW-INTENSITY BROADBAND AND LOW-INTENSITY PULSED ULTRASOUND ON THE AMPUTATIONAL MODEL OF WOUND

**DOI:** 10.64898/2026.04.09.717366

**Authors:** Zaporozhan Valerii, Volokh Karl, Marchenko Olexandr, Godlevsky Leonid, Pervak Mykhailo, Nitochko Oleg

**Author notes:** **Address all correspondence to:** Godlevsky Leonid, Odesa National Medical University, 2, Valikhovsky Lane, Odesa – 65082, Ukraine.

## Abstract

**Background and aim:** Trauma healing with low-intensity ultrasound is effective for different types of injuries affecting both soft tissues and bones. The work aimed to disclose the healing potential of a new type of ultrasound, ultra-wideband low-intensity mechanical waves (UMUS), and to compare its effects with those of low-intensity pulsed ultrasound (LIPUS) in a model of trauma.

**Material and methods:** The work was performed on 2-to 3-month-old male Wistar rats. The model of tail amputation was created, and a transducer emitting UMUS (1-7 MHz, 0.22 mW/cm2) was applied daily for 10 days to the surface of the trauma site in animals that were timely immobilized. LIPUS (1.5 mHz, 30.0 mW/cm2) was used in a separate group of animals. Sham-stimulated rats were used as a control. The intensity of collagen expression in the subdermal tissue was assessed in van Gieson-stained sections, whereas in the UMUS group, expression of CD31, CD34, VEGF, and Ki67 was analyzed.

**Results:** Starting on the 20th day after trauma, UMUS-treated animals demonstrated a statistically significant decrease in the surface area of the traumatic zone compared to the control, whereas LIPUS-treated rats showed this difference on the 30th day of observation. Starting from the 30th day, a significantly greater reduction in the surface of trauma was observed in UMUS, with complete closure achieved in 6 out of 9 rats (P=0.019 vs control), whereas in LIPUS-treated animals, a similar result was observed in 2 out of 8 rats (P>0.05). In UMUS-treated rats, heightened expression of collagen in animals with LIPUS exceeded control data by 7.84% (P=0.034), while the expression in rats with UMUS exceeded data in LIPUS-treated rats by 14.71% (P=0.013). Increased expression of CD31, CD34, VEGF, and Ki67 was observed in UMUS-treated rats.

**Conclusions:** UMUS treatment accelerated healing and reduced wound size, and increased the expression of collagen, CD31, VEGF, CD34, and Ki67, supporting angiogenesis and collagen formation. Effects are more pronounced compared to LIPUS treatment.

**Graphical abstract:** 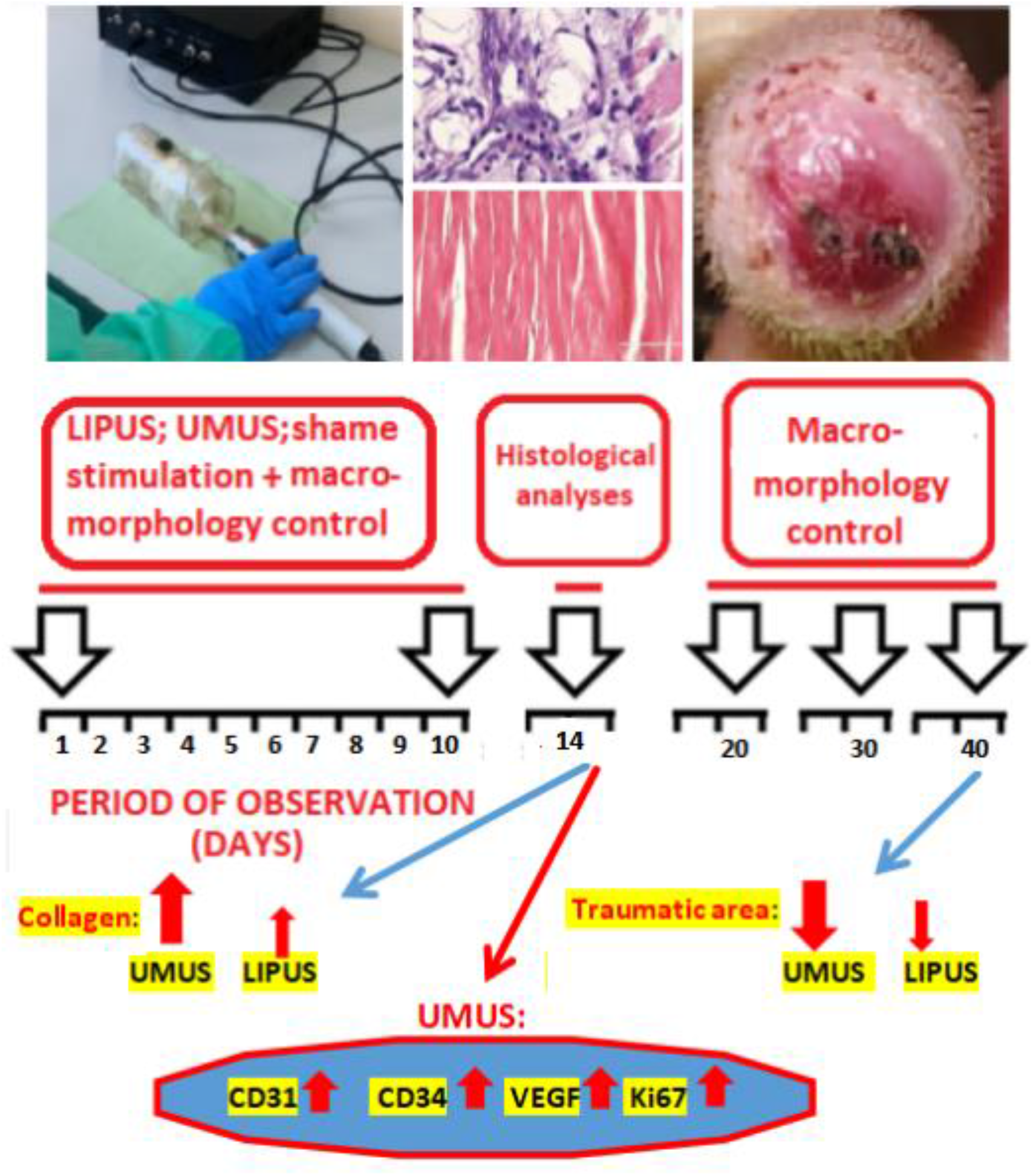

## Introduction

Low-intensity pulsed ultrasound (LIPUS) has been shown to be an effective modulator of numerous diseases, due to its well-established anti-inflammatory effects and stimulation of tissue regeneration [20, 25]. Such fundamental action is initiated by primary excitation of mechanoreceptors, leading to the involvement of numerous intracellular signaling pathways [19].

It is well known that LIPUS promotes proliferation and accelerates healing in soft tissues, such as skin and tendons, as shown by clinical data [16]. These effects occur by activating angiogenesis through increased expression of Vascular Endothelial Growth Factor (VEGF) [33]. The newly formed microvessels boost perfusion of inflamed tissues, helping to restore homeostasis in the wound zone. Activation of mesenchymal stem cells by LIPUS further aids this process [32]. Additionally, LIPUS stimulates fibroblasts and collagen synthesis via the phosphatidylinositol-3-kinase/serine/threonine kinase (PI3K/Akt) and Mitogen-Activated Protein Kinase/Extracellular Signal-Regulated Kinase (MAPK/ERK) pathways, which initiate cell mitosis and the formation of the extracellular matrix. These effects are observed in soft tissues, bones, and cartilage.

LIPUS restricts inflammation, exerts antioxidant effects by modulating endogenous pro- and anti-inflammatory cytokines, and promotes healing [18, 26]. LIPUS also alleviates inflammation via modulation of the nuclear factor-κB (NF-κB) signaling pathway [19], and exosomes - connected mechanisms as well [17]. Via induction, mesenchymal cells LIPUS induced fibroblast activity through the Rho/ROCK pathway and p38 MAPK signaling, leading to faster wound closure [29].

Last time, the ability to prevent neurodegeneration, restrict apoptosis, and improve cognitive functions opened new perspectives in the field of non-invasive neuromodulation and created an alternative approach to the treatment of pharmacologically resistant forms of brain disease [24].

Considering the mechanoreceptive activation role in regenerative medicine, which can be activated by low-intensity ultrasound that avoids thermal effects, an alternative device was created capable of generating ultra-wideband mechanical ultrasound (UMUS) [21]. These low-intensity mechanical waves (UMUS) are characterized by the simultaneous generation of a 1-7 MHz spectrum with pulses duration of 50 μs [14]. It was shown that UMUS induced cytotoxicity and increased mutagenic changes of MCF-7/DDP breast cancer cells *in vitro* [14]. Melanoma B16 cells in vitro also demonstrated cytostatic/cytotoxic pro-apoptotic dynamics induced by UMUS [13]. Peritoneal activity of macrophages increased under UMUS influence in vitro [12]. Additionally, in a mouse model of skin wound (linear dissection), healing was accelerated with 10 sessions of UMUS exposure [22]. Observed effects exceeded those induced by LIPUS. But the molecular mechanisms of the UMUS effects were not disclosed.

Considering the role of various progenitor and stem cell activation in the effects of low-intensity ultrasound [5, 28], the expression of CD34 in the zone of trauma is a possible mechanism for UMUS. The role of progenitor CD34 cells extends beyond endothelial cell formation and includes modulation of cell proliferation and differentiation, cell and tissue architecture, and other functions [25]. Therefore, it is important to also consider the roles of the collagen compartment, angiogenesis, and cell proliferation when investigating these mechanisms.

Hence, the research aimed to investigate the effects of UMUS and LIPUS on the time-course of healing of specific trauma - amputation of rat tail, with the emphasis on collagen production, and to investigate the expression of specific markers CD34, Ki67, CD31, VEGF, as contributors to UMUS effects.

## 2. Material and Methods

### 2.1. Animals

Experiments were performed on 46 male Wistar rats (two to four months old) with an initial body weight of 200–270 g. Animals were kept in standard conditions (constant temperature 23 °C, relative humidity 60 %, 12 h dark/light cycles; standard diet and tap water were given ad libitum) and were acclimatized to laboratory conditions for at least 7 days before the experiment. All experiments were carried out in accordance with the National Institutes of Health Guidelines for the care and use of laboratory animals and the European Council Directive on the Care and Use of Laboratory Animals (86/609/EEC). The experiments were approved by the Odesa National Medical. University Bioethics Committee (UBC) (approval No. 5 dated 11/01/2023) before the study.

### 2.2. Model and macromorphology dynamics measurement

A transverse section of the tail with a scalpel at a depth of 4.0-5.0 cm from the proximal end of the tail was performed under local lidocaine (2%) anesthesia. Chlorhexidine was used locally to prevent infection. Rats were kept in individual plastic boxes, and daily frontal photos of the traumatic surface were taken, with subsequent measurements using ImageJ software. Results of measurements were expressed in mm^2^. Criteria for including cases as having complete healing are presented in Fig. 1.

**Fig. 1.**
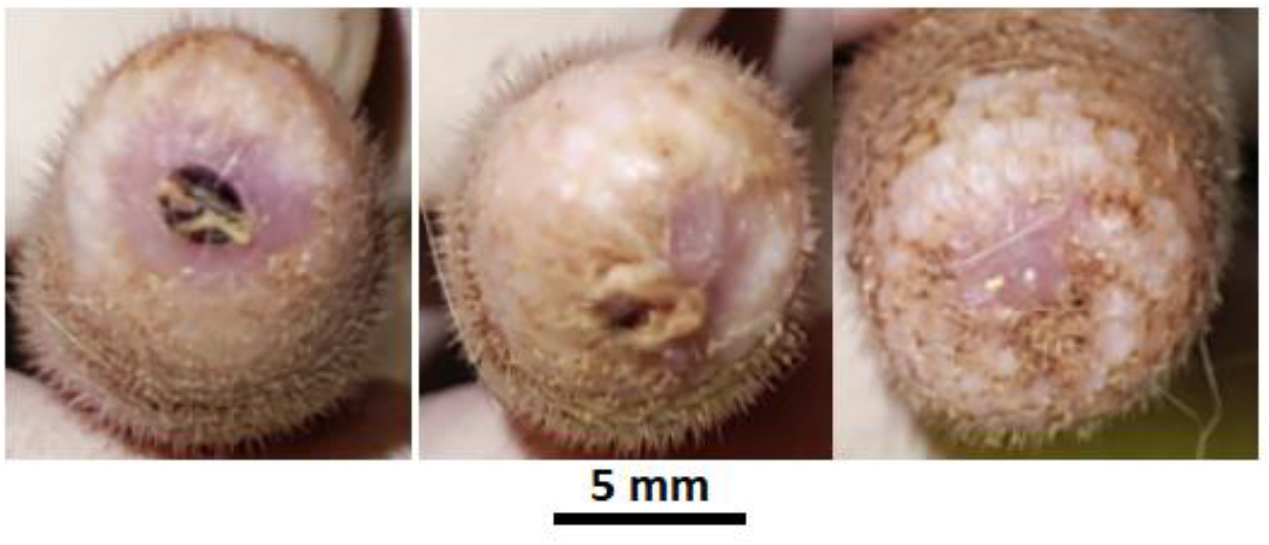
Representative samples of amputation trauma surface assessment indicated as “not complete” (left image) and “complete” healing (subsequent two images).

**Fig. 1.**
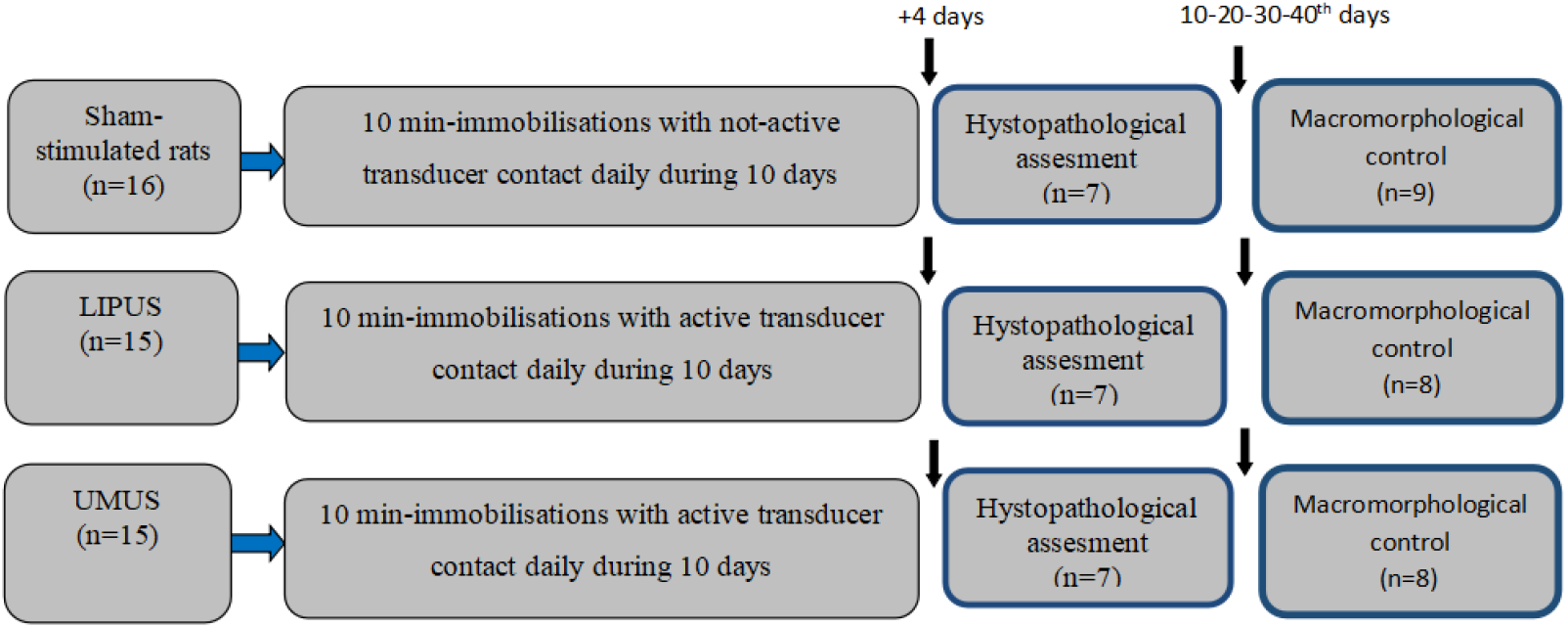
Design of investigation.

Notes from left to right: first photo square not covered with epithelial tissue, the surface of the wound is equal to 19.0 mm^2^, and is considered not completely healed. The next photo shows a square covered with netting measuring 0.6 mm^2^. This is considered complete healing because the minimal area for merged tissue is set at 2.0 mm^2^. The last photo demonstrates complete treatment, with the stump surface covered with epithelium and signs of wool growth.

### 2.2. Design of investigation

Three groups of rats were formed and observed for 40 days, starting the day after the trauma (Fig. 1).

During traumatic surface exposure to the transducer, animals were immobilized in a plastic tube, and tight, highly controlled contact with the transducer surface was maintained for 10 minutes. The ultrasonic gel Aqua Ultra Basic UBQ Ultragel (Hungary) was used as a conductive medium. In the control group, the same procedure was performed, but the ultrasound generator was not turned on. In total, 10 sessions were conducted in each group.

Histological tissue was obtained on the 14^th^ day of observation by sectioning 1.5-2.0 cm of the tail stump under local anesthesia in seven rats, which were randomized using a blinded method within each group. The remaining rats were macromorphologically monitored until the end of the observation period.

### 2.3. Characteristic of UMUS and LIPUS waves

An SDG 2082 X generator from Siglent was used as a source of electrical signals for excitation of the ultrasonic transducer. Ultra-wideband conversion of generator signals into ultrasonic vibrations in the frequency band 1-7 MHz, intensity of 0.22 mW/cm^2^, and duration of 50 μs was carried out by a 15-mode piezoceramic transducer with overlapping vibration modes and polarization normal to the radiating surface [14]. The transducer has a diameter of 20 mm and a maximum thickness of 3 mm. Similarly, LIPUS was got as a result of conversion to 1,5 MHz, power of 30.0 mW/cm^2^ and pulse duration 200 μs [12-14].

### 2.4. Tissue samples preparation and histological analysis

#### 2.4.1. van-Gieson staining

For van-Gieson staining, tissues were embedded in molten paraffin wax at 60°C, and sections were cut at 4 microns on a rotary microtome. All sections were dried for 30 min at 60°C in readiness for staining [2].

Digital micrographs of Van Gieson-stained sections were analyzed using Adobe Photoshop CS5 Extended, version 12.1 x64 (Adobe Systems Incorporated, San Jose, CA, USA). Identical parameters for red color-range selection were applied to all images using the Color Range tool. For each image, the proportion of pixels assigned to the red color range was calculated relative to the total analyzed field area.

#### 2.4.2. Immunohistochemical staining

Tail tissues were fixed in 10% formaldehyde, washed in tap water (running water for 10 hours), and kept in increasing alcohol series for the water recovery procedure followed by 30 min reeping in xylol. After that, tissues were embedded in paraffin, and 5 μm-thick sections were cut from the blocks and placed on slides coated with poly-L-lysine for immunohistochemistry analysis.

The avidin-biotin peroxidase complex method was used to determine differences in Ki67, CD31, VEGF, and CD34 expression in brain tissue, and expression was performed in accordance with the earlier-described method [23]. For the incubation the Abcam antibodies were used: CD31 (EPR17260-254 / ab213175); VEGF (ab100786); Ki67 [SP6] (ab16667), and for CD34 - AF4117 (R&D Systems). Results of immunohistochemical staining were evaluated based on the presence of specific brown-colored morphological structures, as confirmed by the negative control [7].

### 2.5. Statistics

The SPSS program (SPSS for Windows, SPSS Inc., Chicago, IL, USA, version 21.0) was used to perform statistical analysis. Data are given as mean values ± standard deviation of the mean (SD) and presented as violin diagrams with open access Statistics Kingdom [https://www.statskingdom.com/violin-plot-maker.html]. Comparison of macromorphological data between groups was analyzed using ANOVA and Tukey HSD post hoc tests, and the “z” criterion for proportions was used to determine rats with complete wound healing. For comparing the number of red points in the collagen differences measurement, the Kruskal-Wallis test with Dunn’s post hoc test was appropriate. Differences were considered significant at *\**P< 0.05.

## Results

On the 20^th^ day after trauma, a significant difference in trauma area was observed between groups (F=16.162; df=2,24; P=0.000; Fig. 1). The trauma area in UMUS-treated rats was reduced by 16.5% compared to LIPUS-treated rats (P=0.006), while LIPUS-treated animals did not show a significant difference from the control group (P=0.1173).

One month after trauma, significant differences between groups persisted (F=18.378; df=2,22; P=0.000). During this period, the trauma area in the LIPUS-treated group decreased by 44.9% (P=0.0076), while in UMUS-treated animals it was 61.1% smaller than in the LIPUS-treated group (P=0.042). These group differences continued at day 40 (F=13.542; df=2,22; P=0.0001; Fig. 1). At this point, the trauma area in LIPUS-treated animals was 45.4% less than in controls (P=0.045), and UMUS-treated animals (5 rats) again showed a significantly smaller trauma area than LIPUS-treated animals (P=0.0428). Moreover, closure was observed in 6 of 9 UMUS-treated rats (z=2.339; P=0.019), compared with 2 of 8 LIPUS-treated rats (z=0.756; P=0.450).

In van-Gieson-stained slices, characteristic pink - red collagen fibers were identified with yellow bright cytoplasm and brown dark colors of nuclei of fibrocytes, and muscle cells. In the control group, thin fibers not organized into bundles were observed (Fig. 4A). In rats treated with LIPUS, regular bundles of collagen fibers were present (Fig. 4B). In rats treated with UMUS, intensely colored, thick bundles of collagen fibers, regularly oriented, were observed (Fig. 4C). Quantification of red pixels revealed statistical differences between groups at H=21.8959; P=0.000. Namely, animals with LIPUS exceeded control data by 7.84% (P=0.034), and rats with UMUS exceeded data in LIPUS-treated rats by 14.71% (P=0.013) (Fig. 4).

**Fig.3.**
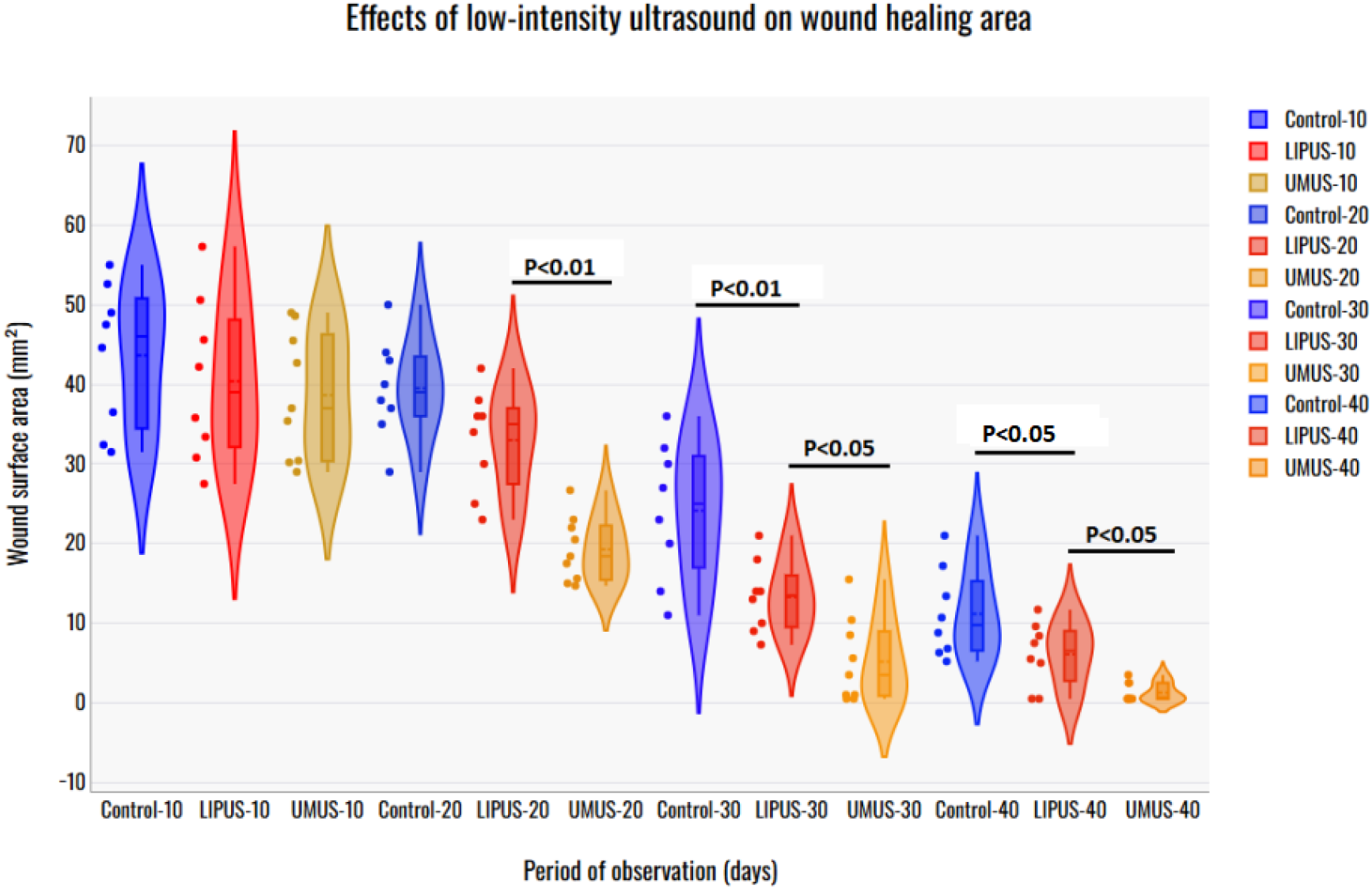
Dynamics of the traumatic surface reduction after a 10-day course of treatment with low-intensity ultrasound. Notes: The number of days from the first treatment session is marked for each observation group. ANOVA+Tukey HSD was used for wound surface quantitative comparison.

**Fig.4.**
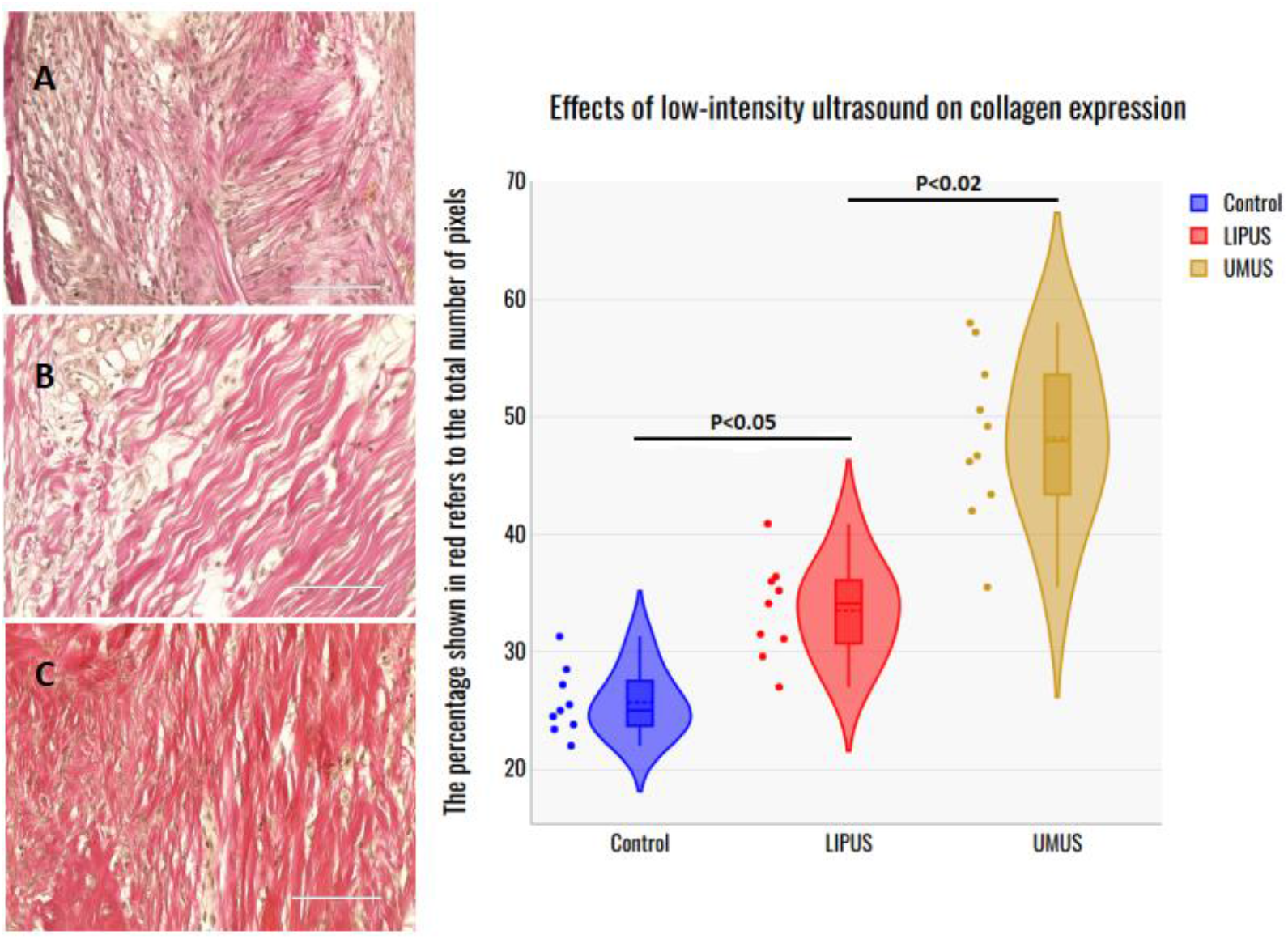
Quantification of collagen expression in van-Gieson-stained slices in sub-dermal tissue by the number of red pixels in Van Gieson-stained samples. (14th day from the moment of treatment start). A-Control; B-LIPUS-, and C-UMUS treatment. Kruskal-Wallis and Dunn’s post hoc test were appropriate.

Immunohistochemical staining of the sub-dermal zone revealed thick bundles of collagen surrounded by numerous cells, presumably fibrocytes (blue-stained cells), that are partially marked with Ki-67 as fibroblasts in 2-7 mm depth from the surface of trauma (Fig. 5, A). Characteristic circular structures were identified in bone tissues and in subdermal neighboring tissue with VEGF staining (Fig. 5, B). Similar circular structures were observed in sub-dermal tissue with CD31 staining (Fig. 5C) and with CD34 staining in both trabecular bones (Fig.5, D).

**Fig. 5.**
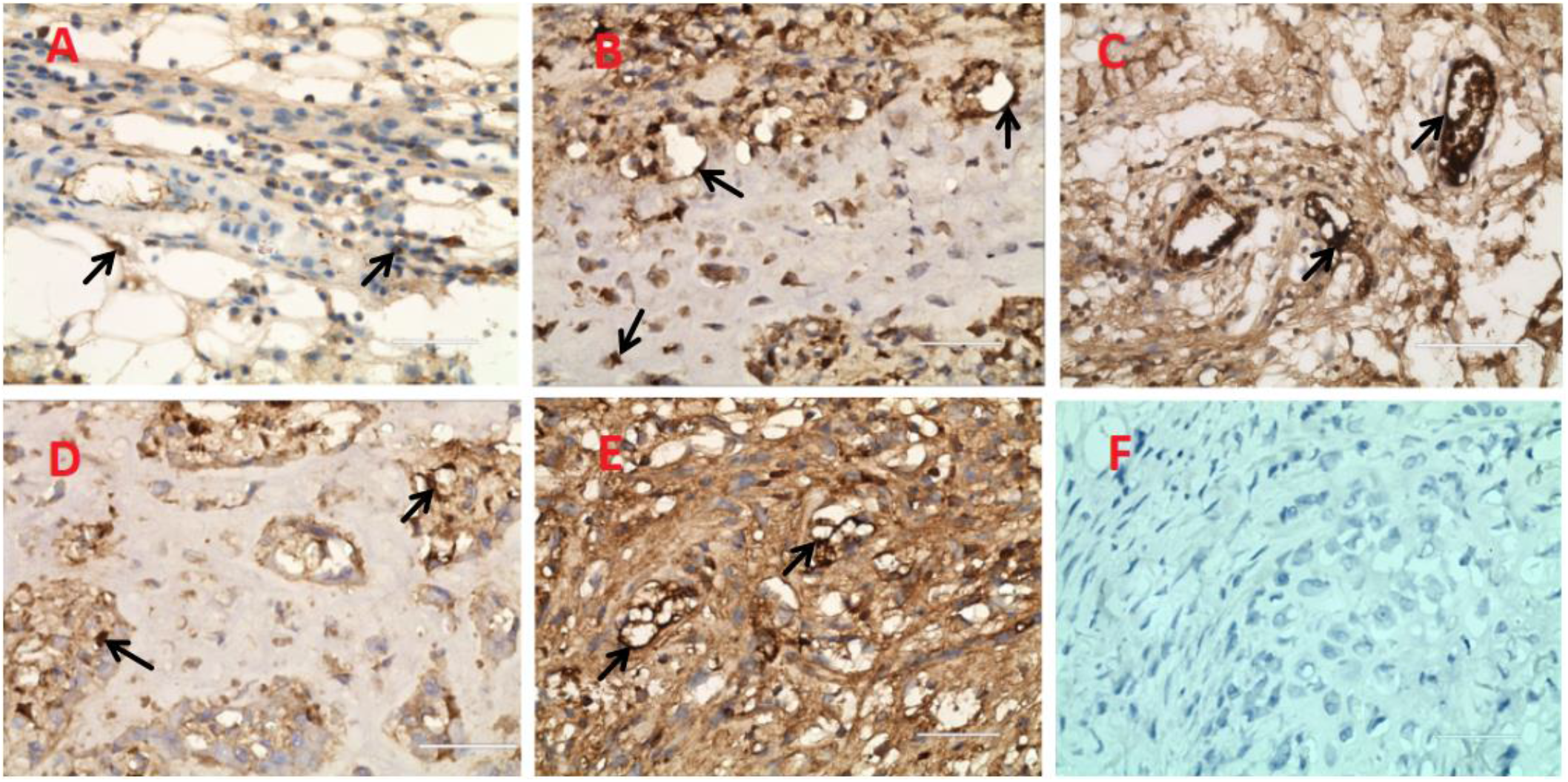
Immunohistochemical tissue staining in rats treated with UMUS on the 14th day from the moment of treatment start. A-Ki-67 in sub-dermal tissue marked with arrows; along collagen thick bunches, fibrocytes are gathered, stained with blue color; some of them are marked as blast forms (arrows); B - VEGF stained structures in both bones and neighboring sub-dermal tissue. C - CD31 in sub-dermal tissue (arrows); D and E - CD34 in bone and sub-dermal tissue, respectively; F-negative control. Calibrate at 100 mcm.

## Discussion

UMUS effects achieved a significant level of shortage of the traumatic area on the 20^th^ day of the posttraumatic period, while LIPUS induced a significant difference on the 30^th^ day when compared with the control data. The most effective UMUS healing was characterized by a significant reduction in the surface area of traumatic amputation starting on the 20th day of observation, compared with LIPUS-treated animals, with the consequent maintenance of this difference until the 40^th^ day of the experiment. Additionally, on the 40th day, complete closure of the traumatic surface was observed in 6 out of 9 rats treated with UMUS.

Clear stimulation of collagen synthesis in the traumatic zone caused by UMUS might be conditioned by the specificity of collagen fibers/collagen-containing tendons, as it plays a fundamental role in tail functionality. Thus, rat tail tendons are composed almost entirely of highly aligned, dense Type I collagen fibers, whereas rat skin contains a mixture of Type I and Type III collagen [1, 27]. It should be noted that skin-induced trauma in mice was also effectively healed with UMUS treatment, with healing effectiveness higher than with LIPUS treatment [22]. But the differences and dynamics of wound healing were less pronounced than in our data. Such a difference might be explained by both the exploration of anesthesia in mouse experiments and the aforementioned differences in collagen composition between skin and tail tissue.

Hence, skin trauma might be less prone to demonstrating collagen synthesis than tail tissues. That is why, although Ki67 is not a specific marker for fibrocytes, the mentioned peculiarity, along with the time schedule of collagen genesis (maximal after the 10^th^ day after trauma), favors their contribution to the observed manifestations. In this context, it is important to mention that CD34 is widely employed in diagnostic breast pathology as an endothelial marker, but it is also expressed by a population of non-endothelial stromal fibroblastic cells [20, 25]. Hence, considering the high level of collagen in tail tissue, the time-course of active fibroblasts’ involvement in collagen production, established Ki67 reactivity in terms of 10-15 days after trauma, favors the presence of numerous fibroblasts in the zone of high accumulation of collagen in 9-14 days after skin trauma [9]. Comparatively less pronounced LIPUS effects might also be explained by its stimulatory effects on collagen Type II, as shown for extracellular matrices during chondrogenesis [28].

The next mechanism of wound healing, well known for LIPUS action, is its anti-inflammatory effects [17, 19]. In this context, activation of macrophages induced with UMUS is also favored, as UMUS exhibits similar activity [12]. Gained data on the presence of CD31 -platelet endothelial cell adhesion molecule-1 (PECAM-1) and VEGF expression favors increased angiogenesis caused by UMUS. Indeed, LIPUS stimulates the expression of CD31, a marker of endothelial cells in the endometrium, and VEGF, a marker of angiogenesis [8, 16]. VEGF, induced with LIPUS, is in charge of angiogenesis promotion in soft tissues [33]. Besides, the expression of the progenitor cell marker CD34 is also increased [16].

Considering the net mechanisms that might underlie the observed effects of UMUS, it should be noted that low-intensity ultrasound affects mechanoreceptors, which are recognized as the first step in the precipitation of subsequent positive therapeutic effects. Thus, mechanical microvibration disturbed tissular piezoelectric materials and generated an electric signal that influenced cell properties, including differentiation and proliferation, functional shift in excitatory cells, and regenerative processes [4, 10, 11]. The important consequence of mechano-electric transduction is the involvement of ionic channels and a change of ionic environment of the excitatory membrane with activation of intracellular signaling pathways after the appearance of ionized calcium in the intracellular space [15. 30]. It should be noted that integrins, especially abundant in bone tissue, function as mechanoreceptors for low-intensity ultrasound [31]. Caveolae, as a target for ultrasound waves, are presented as small pits on the membrane measuring 50-100 nm in size that, under mechanical stress, are flattened and elaborate their content intracellularly, serving as an initial informative signal for intracellular metabolism [6, 15]. It should be stressed that”wade-band” ultrasound is able under certain conditions to maximize the high-frequency components and their nonlinear interactions (harmonics), it can generate localized mechanical stress gradients that match the dimensions of caveolae [3]. This would allow for “nanomechanical” stimulation that LIPUS simply cannot achieve due to its long, stable wavelengths.

Finally, it should be stressed that tail amputation trauma in rats is a new and excellent model for wound healing, as it includes soft tissues and bones, a substantial amount of collagen, and treatment does need anesthesia and might be performed with minimal stress to animals with very reliable contact with the surface of ultrasound emission. The last position is of great significance for UMUS, as the emitted energy is less than that of LIPUS.

## Conclusions

1. UMUS accelerated healing and wound reduction more than LIPUS, with differences observed by day 20.
2. UMUS more strongly activated collagen formation than LIPUS.
3. UMUS increased expression of CD31, VEGF, CD34, and Ki67, supporting angiogenesis and collagen formation.
4. The rat tail amputation model is reliable for studying collagen-based regeneration and allows precise ultrasound application.

## Acknowledgments

The authors express their gratitude to Dr. Natalya Bukreeva, chief of the molecular-genetic laboratory, assistant professor Kseniya Pribolovets, and students Dmytro Arabadji, Illya Shcheglov, and Vladimir Bidnyuk for the technical support of experimental observations, to assistant professor Sergyi Marchenko for the analysis of data, all from Odesa National Medical University. Also, thanks to Dr. Olexandr Shcheglov for engineering support (“Techno Med Ukraine”, Kyiv, Ukraine)

